# Differential network analysis of human tissue interactomes highlights tissue-selective processes and genetic disorder genes

**DOI:** 10.1101/612143

**Authors:** Omer Basha, Chanan M. Argov, Raviv Artzy, Yazeed Zoabi, Idan Hekselman, Liad Alfandari, Vered Chalifa-Caspi, Esti Yeger-Lotem

## Abstract

**Motivation:** Differential network analysis, designed to highlight interaction changes between conditions, is an important paradigm in network biology. However, network analysis methods have been typically designed to compare between few conditions, were rarely applied to protein interaction networks (interactomes). Moreover, large-scale benchmarks for their evaluation have been lacking.

**Results:** Here, we assess five network analysis methods by applying them to 34 human tissues interactomes. For this, we created a manually-curated benchmark of 6,499 tissue-specific, gene ontology biological processes, and analyzed the ability of each method to expose these tissue-process associations. The four differential network analysis methods outperformed the non-differential, expression-based method (AUCs of 0.82-0.9 versus 0.69, respectively). We then created another benchmark, of 1,527 tissue-specific disease cases, and analyzed the ability of differential network analysis methods to highlight additional disease-related genes. Compared to a non-differential subnetworks surrounding a known disease-causing gene, the extremely-differential subnetwork (top 1%) was significantly enriched for additional disease-causing genes in 18.6% of the cases (p≤10e-3). In 5/10 tissues tested, including Muscle, nerve and heart tissues (p = 2.54E-05, 2.71E-04, 3.63E-19), such enrichments were highly significant.

**Summary:** Altogether, our study demonstrates that differential network analysis of human tissue interactomes is a powerful tool for highlighting processes and genes with tissue-selective functionality and clinical impact. Moreover, it offers expansive manually-curated datasets of tissue-selective processes and diseases that could serve for benchmark and for analyses in many other studies.

## INTRODUCTION

Knowledge of the molecular compositions of human tissues and cells has grown immensely in recent years, owing to large-scale omics projects such as the Human Protein Atlas (Uhlen *et al.*, 2015), the Genotype-Tissue expression (GTEx Consortium, 2017) and, most recently, the Human Cell Atlas (Regev *et al.*, 2017). To better understand the functions of tissues and cells, continuous efforts were invested in charting their underlying molecular interactions, particularly among human proteins (Luck *et al.*, 2017). To date, over 360,000 protein-protein interactions (PPIs) among over 18,000 proteins have been detected experimentally and recorded in public databases. Integrating between these PPIs and tissue expression profiles enabled the construction of network models (interactomes) of human tissues (Yeger-Lotem and Sharan, 2015). Compared to a generic interactome, tissue interactomes were shown to be useful for identifying tissue-specific protein functions (Greene *et al.*, 2015), and for prioritizing candidate disease-associated genes (Greene *et al.*, 2015; Magger *et al.*, 2012; Kitsak *et al.*, 2016).

The molecular profiling of human tissues revealed that distinct tissues share most of their expressed genes with all other tissues (GTEx Consortium, 2017), and, consequently, much of their underlying interactions (Barshir *et al.*, 2014). This called for analyses that will be able to explain tissue-specific phenotypes. Several such methods focused on the identification of tissue-specific genes and interactions, used for example to illuminate the molecular mechanisms underlying tissue-selective disease manifestations (e.g., (Barshir *et al.*, 2014), as well as other applications, as reviewed in (Yao *et al.*, 2018). Also unique and often ignored, were genes and interactions that were down-regulated in a certain tissue relative to other tissues. Such down-regulated genes were recently associated with the predisposition of tissues to hereditary diseases (Barshir *et al.*, 2018).

Differential network (DN) analysis is a powerful paradigm that highlights genes and interactions that are either up-or down-regulated in a specific context (Ideker and Krogan, 2012). Accordingly, genes and interactions that are shared among contexts are downplayed. Many of them could be carrying important house-keeping functions, however, they have limited ability to illuminate the mechanisms underlying context-specific phenotypes. In contrast, these mechanisms can be illuminated by focusing on the altered genes and interactions (Bandyopadhyay *et al.*, 2010). DN methods were applied to multiple types of molecular network models, including genetic interaction networks (Bandyopadhyay *et al.*, 2010), co-expression networks (Ma *et al.*, 2014; Pierson *et al.*, 2015), PPI networks (Basha, Barshir, *et al.*, 2017), and regulatory networks (Marbach *et al.*, 2016). Computationally, DN methods focused on network nodes (e.g., (Sonawane *et al.*, 2017), (Basha, Barshir, *et al.*, 2017)), network interactions (e.g., (Basha, Shpringer, *et al.*, 2017)), or network features such as node connectivity (e.g., (Goenawan *et al.*, 2016)). DN methods proved useful for prediction of cancer genes (Islam *et al.*, 2013; Warsow *et al.*, 2013), drug targets (Zickenrott *et al.*, 2016), and plants’ stress response genes (Ma *et al.*, 2014). Several DN methods were implemented as open tools (e.g., (Gill *et al.*, 2010; Landeghem *et al.*, 2016; Gambardella *et al.*, 2013; Ha *et al.*, 2014)). In particular, we developed a DN method for tissue interactomes and made it available through the DifferentialNet database (Basha, Shpringer, *et al.*, 2017). Specifically, each PPI within a tissue interactome was associated with a score, reflecting the up-or down-regulation of the interacting proteins relative to other tissues. Yet, a rigorous assessment of DN methods applied to PPI networks in general and to tissue interactomes in particular has been lacking, mostly for lack of suitable, large-scale benchmarks.

Here we assess the performance of a non-differential method and four node-based and interaction-based DN methods, which we applied to analyze 34 human tissue interactomes. To create these interactomes, we integrated the rich dataset of RNA-sequenced human tissue profiles of gathered by GTEx (GTEx Consortium, 2017) with data of over 333,000 experimentally detected PPIs. We implemented the five methods and tested their ability to capture tissue-specific features. For this, we created a manually curated dataset associating 6,499 gene ontology (GO) biological processes to their relevant human tissues. DN methods performed better than a non-differential method in accurately highlighting tissue-specific processes. We also evaluated node-based versus interaction-based methods.

Next, we applied DN methods to illuminate the enigmatic tissue-selective manifestation of hereditary diseases; though caused by aberration in genes that are typically expressed across many tissues, hereditary diseases tend to manifest in few, selected tissues (Barshir *et al.*, 2014, 2018). Specifically, we tested whether the altered sub-network that surrounds disease-causing genes in their respective disease-tissues is enriched for additional disease-relevant genes. For this, we manually associated 1,185 tissue-selective hereditary disorders with their clinically manifesting tissues. In 18.6% of the cases, enrichment of the top 1% altered subnetwork was highly statistically significant, a fraction much higher than expected by chance based on permutation testing (p≤10e-3). Enrichment was specifically high in muscle, nerve and heart (p = 2.54E-05, 2.71E-04, 3.63E-19) tissues. Thus, DN analysis of tissue interactomes offers a powerful filtering approach for identifying tissue-selective physiological processes and disease genes.

## RESULTS

### Constructing tissue interactomes

We started by creating an up-to-date version of the human interactome by gathering experimentally-detected PPIs from multiple databases (see Methods). This resulted in a generic interactome containing 18,542 proteins and 333,745 interactions that were detected by well-established detection methods (Basha *et al.*, 2015). To construct interactome models for different human tissues, we used RNA-sequenced profiles pertaining to 51 tissues that were gathered by GTEx (GTEx Consortium, 2017). To avoid a bias toward brain regions, brain sub-regions were further merged into basal ganglia, cerebellum, and ‘other’ (as in (Paulson *et al.*, 2017)), resulting in 40 different tissues (see Methods). Profiles were normalized as described elsewhere (Basha, Barshir, *et al.*, 2017). Our next step was to integrate the generic human interactome with the tissue expression profiles to create tissue interactomes.

We implemented five different methods, including three node based and two edge-based methods (see Methods and Table 1). The first was a non-differential, node-based method, denoted *expr_n,* where each gene in a tissue interactome was assigned with a weight that reflected its expression level in that tissue. Accordingly, genes with high, uniform levels across tissues will be assigned high weights in all tissue interactomes. Next, we implemented two differential node-based methods. In the first, denoted *pref_n,* each gene in a tissue interactome was assigned with its preferential expression in that tissue, as computed by (Sonawane *et al.*, 2017). In the second, denoted d*iff_n*, each gene in a tissue interactome was assigned with the difference between its expression level in that tissue and its median expression level across all tissues. In both methods, genes with high, uniform levels across tissues will be assigned weights close to zero. Based on the two differential node-based methods, we implemented two differential interaction-based methods. In the first, denoted *pref_i*, interaction weight reflected the summed preferential expression of the two pair-mates in that tissue. In the second, denoted *diff_i*, interaction weight was set to the summed expression levels of the two pair mates in that tissue, minus the median summed weight across tissues (Basha, Shpringer, *et al.*, 2017).

**Table 1.** The five different schemes to create tissue specific interactomes.

| Method name | Weighted Entities | Weighting method | Differential |
| --- | --- | --- | --- |
| *expr_n* | Nodes | Gene Expression | Non-differential |
| *pref_n* | Nodes | Preferential expression, developed by (Sonawane *et al.*, 2017) | Differential |
| *diff_n* | Nodes | Computed as the difference between the expression level of a node in a tissue, and its median expression level across tissues. | Differential |
| *pref_i* | Interactions | Computed as the sum of the preferential expression of the interacting nodes, based on (Sonawane *et al.*, 2017). | Differential |
| *diff_i* | Interactions | Computed as the difference between the summed expression levels of the interacting nodes in the tissue and their median summed expression across tissues (see methods). Developed by (Basha, Shpringer, *et al.*, 2017). | Differential |

We created tissue interactomes by applying each method to the generic human interactome. Since some genes were not found to be expressed in any tissue, the resulting generic interactome included 16,177 nodes and 257,200 interactions. Notably, all tissue interactomes contained the same nodes and edges, and differed from each other only in the weights assigned to their nodes or interactions. In general, the different methods resulted in normally-distributed weights per tissue (Fig. S1).

### Assessing the methods’ ability to highlight tissue-specific processes

We aimed to evaluate rigorously and at large-scale whether the different methods helped illuminate the distinct features of each tissue. For this, we created a gold set of tissue-specific GO biological process terms. Our preliminary dataset included GO terms whose description contained tissue-related keywords (e.g., ‘adipo’, ‘fat’), as well as previously published tissue-associated GO terms (Greene *et al.*, 2015), which we matched with the tissues profiled by GTEx. We then manually checked each GO term to verify its tissue associations (Fig. 1A). In total, we associated 6,499 terms to 48 tissues through 7,718 associations (Fig. 1B and Table S1).

**Fig. 1.**
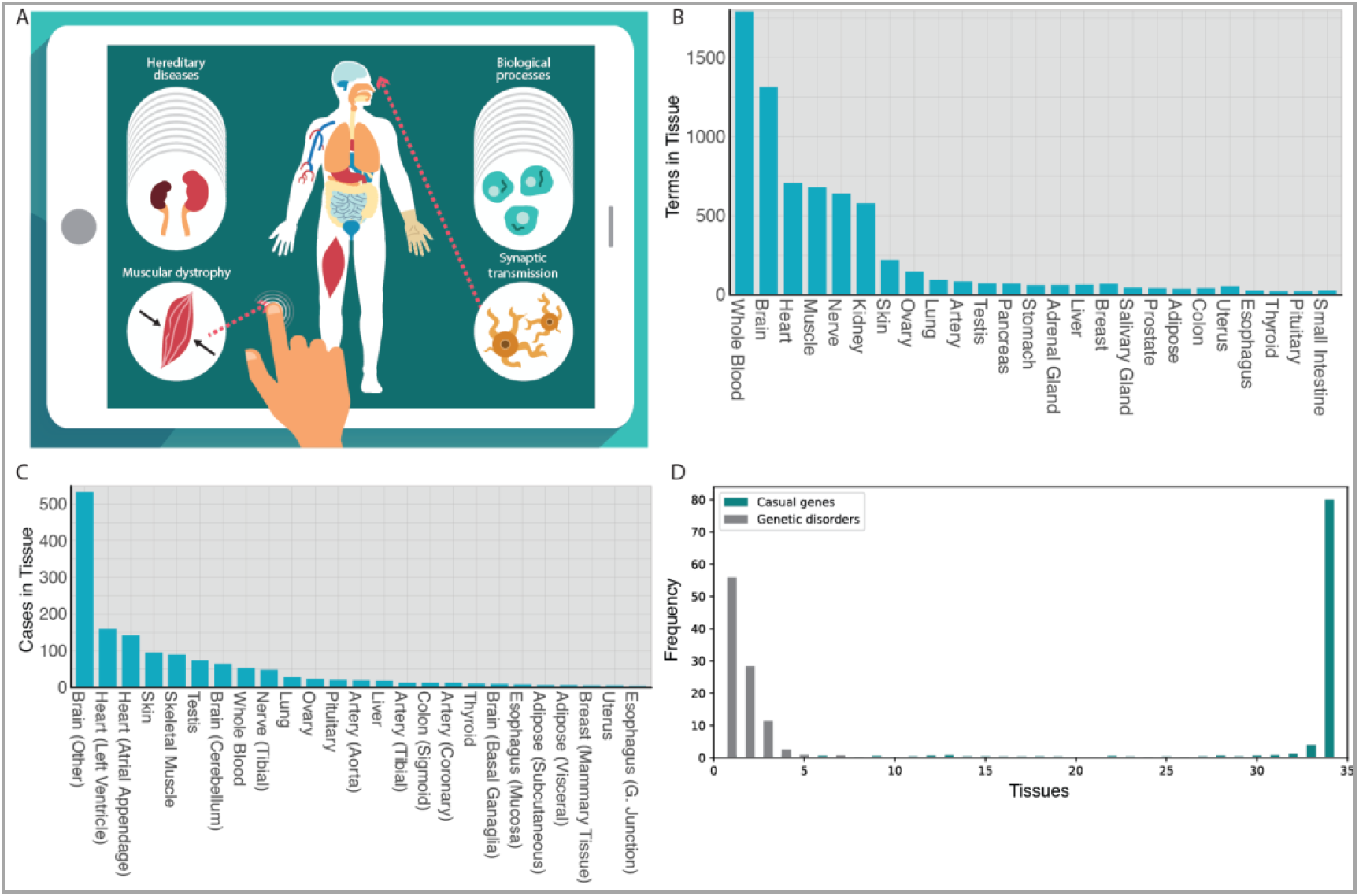
Manually-curated datasets of tissue-specific biological processes and hereditary disorders. A. We manually associated 6,499 GO biological process terms with their relevant tissues in GTEx, and 1,185 hereditary disorders with their disorder-manifesting tissue. For example, the process of synaptic transmission was associated with the brain (right), and muscular dystrophy was associated with muscle (left). B. The number of GO biological process terms that were associated with each tissue. C. The number of hereditary disorders cases (disorder & causal gene) that were associated with each tissue. D. The number of tissues that clinically manifest a disease (grey) or express a disease-causing gene (turquoise). The tissue-selectivity of the 1,185 hereditary diseases stands in large contrast with the ubiquitous expression of their 852 causal genes. A gene was considered expressed in a tissue if its expression value was ≥ 8 normalized counts.

Given this rich dataset, we checked whether the different tissue interactomes were able to highlight their respective tissue-associated processes. For this, we converted each interactome to a ranked gene list. For node-based methods, ranking was determined by node weights in the respective tissue interactome, such that highly expressed genes ranked at the top. For interaction-based methods, we assigned each node with the median weight of its interactions in the respective tissue interactome. Each ranked gene list was then subjected to GO enrichment analysis by using the GOrilla tool (Eden *et al.*, 2009), which allowed us to test per interactome and method whether the corresponding gene list was enriched for GO terms that were associated with the respective tissue. We then calculated per method the number of tissue interactomes showing accurate enrichments (Fig. 2A). The non-differential approach, *expr_n*, had the smallest fraction of accurately enriched tissues (55%). The DN methods had much higher fractions of accurately enriched tissues (73-91%), with *diff_i* performing best.

**Fig. 2.**
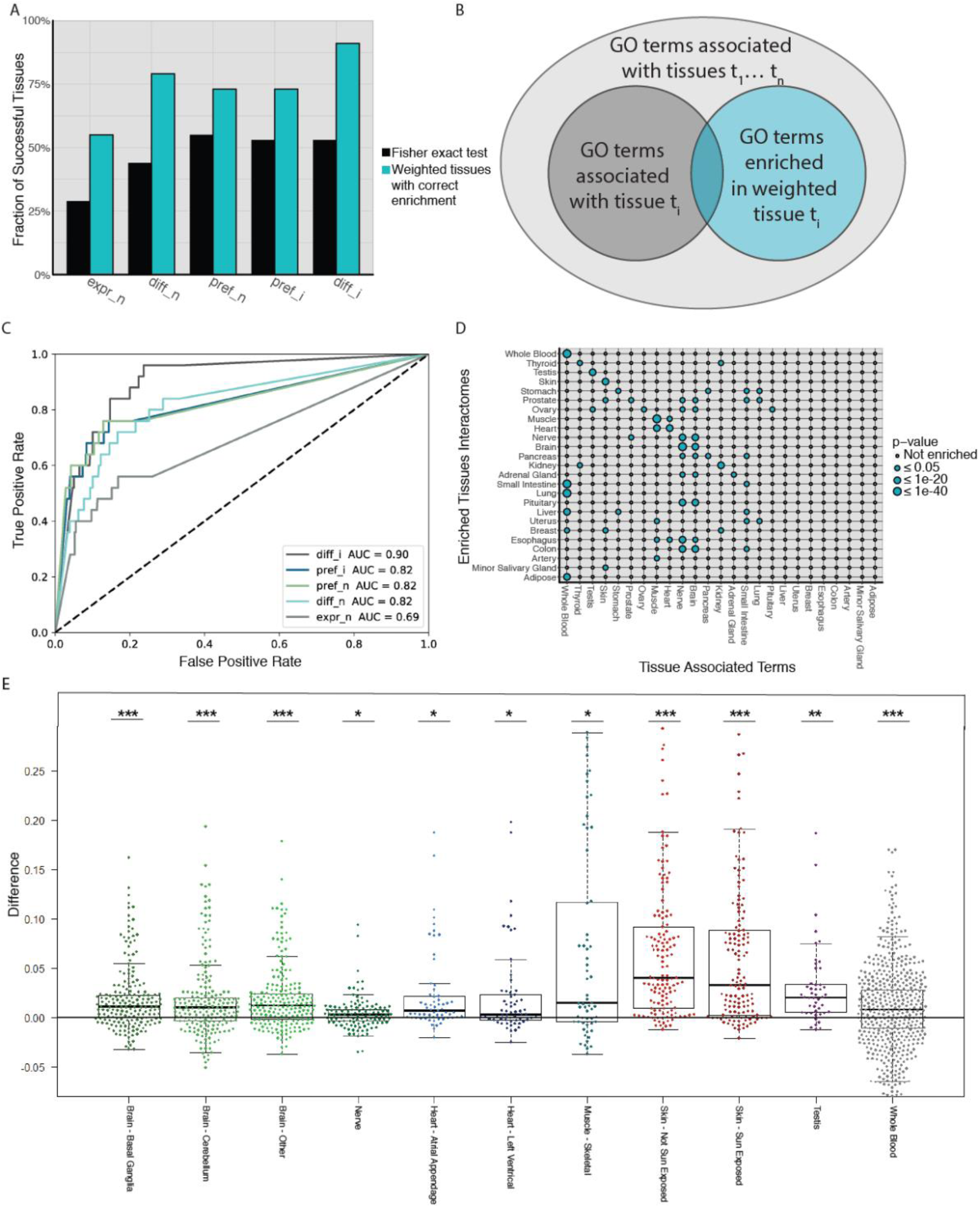
Assessing the ability of weighted tissue interactomes to uncover tissue-specific processes. A. The fraction of weighted tissue interactomes out of the 34 tissue interactomes that were correctly enriched for their respective tissue-associated GO terms (turquois, p≤10E-3), or showed enrichment specificity for these terms (dark color, p≤0.05), per method. B. Illustration of the enrichment specificity test. The number of tissue-associated GO terms that were accurately enriched in the corresponding tissue interactome (overlap area, colored turquoise) was assessed for statistical significance via Fisher exact test. C. A ROC plot showing the prediction accuracy of each method. Differential methods outperformed the non-differential method. D. Visualization of the enrichment specificity test for the *diff_i* approach. Each dot represents the enrichment specificity of the tissue interactome (row) to tissue-associated GO terms (column). Most differential tissue interactomes showed enrichment specificity for their respective terms or for terms of physiologically related tissues. E. Analysis of differential interactions per query gene, for each of the 11 tissues with over 20 query genes. Each dot represents a query gene in the respective tissue. Its value corresponds to the difference between two medians: the median weight of its interactions with proteins sharing its tissue-associated terms, and the median weight of its other interactions. In most query genes, the difference was positive, meaning that interactions involving proteins sharing tissue-associated terms weighted higher. Significance was assessed via paired Wilcoxon test; * p≤0.05, ** p≤1E-5, *** p≤1E-10.

We further assessed the enrichment specificity of tissue interactomes. Explicitly, we tested whether a given tissue interactome was not only enriched for its associated GO terms, but was also more frequently enriched for these terms relative to terms associated with other tissues, by using Fisher exact test (Fig. 2B, see Methods). We then calculated per method the number of tissue interactomes showing significant enrichment specificity (p≤0.05, Fig. 2A). The non-differential method had the lowest fraction of significant tissue interactomes, while *diff_i* and *pref_n* had the highest fractions (Fig. 2A; Table S2). We further compared the enrichment specificity of all methods via a ROC analysis (Fig. 2C). The area under the curve (AUC) was smallest for the non-differential method (AUC=0.69), while all differential methods performed comparably well, especially in the low range of true-positive rate (AUCs of 0.82-0.9). As shown in Fig. 2D for the *diff_i* method, differential tissue interactomes tended to be most enriched for their respective tissue GO terms, or for terms associated with physiologically-related tissues, such as brain and nerve, or skeletal muscle and heart.

Since the previous test relied on gene ranking, we developed an alternative test that focused on interactions, which we applied to 2,332 query genes that were annotated to tissue-specific processes (see Methods). For each query gene in its respective tissue, we tested whether its interactions with proteins sharing its tissue-associated terms weighted significantly more than its other interactions. Interaction weights corresponded to the *diff_i* method, and statistical significance was tested via the Mann-Whitney-U test. This was indeed the case for 376 (16%) query genes (adjusted p≤0.05). To obtain a view per tissue, we collated all query genes associated with the same tissue. Next, we computed, per query gene, its median interaction weights with (i) proteins sharing its tissue-associated terms, and (ii) other proteins, which we inserted into two distinct lists. We focused on tissues with at least 20 query genes, and evaluated the differences between the two lists via a Paired-Wilcoxon test. In all 11 tissues tested, the difference between the two lists was statistically significant (adjusted p≤0.05, Fig 2E). Altogether, these results indicate that differential tissue interactomes are better than a non-differential interactome in highlighting tissue-specific features. This analysis also implies that tissue-specific processes tend to involve differentially up-regulated nodes and interactions.

### Applying the differential approach to reveal genes causal for tissue-specific disorders

Previous studies showed that hereditary disorders tend to manifest in few specific tissues, although their causal genes are present and often expressed throughout the body (Lage *et al.*, 2008; Barshir *et al.*, 2014, 2018). The molecular mechanisms underlying hereditary disorders, and in particular their tissue selective manifestation, remain unclear for most disorders. Here, we tested whether differential tissue interactomes may help illuminate these underlying mechanisms. To enable a large-scale assessment, a goldset of tissue associated diseases was needed, yet none were available. Therefore, we manually associated phenotypic series from the OMIM database with the tissues in which they manifest clinically by reviewing their description in OMIM and other medical references (See Methods). A phenotypic series is a group of genetic diseases that manifest with similar phenotype. We focused on disorders with known causal genes and that were phenotypically related according to OMIM (Amberger *et al.*, 2015)(see Methods, Fig. 1A,C, Table S3). Altogether, we analyzed 1,185 disorders with 1,527 gene cases associated to 26 tissues and 852 causal genes. Notably, while most disorders were highly tissue-specific, most of their causal genes were expressed widely across tissues (Fig. 1D). Thus, it was intriguing to ask whether the differential interactions surrounding causal genes in their respective disease-manifesting tissues could unravel disorder-related mechanisms.

We considered for each causal gene the interactome subnetwork containing its direct and secondary interactions (Fig. 3A, left panel). We focused only on subnetworks that included additional genes causal for a similar disorder. To test the ability of differential tissue interactomes to reveal disorder-related mechanisms, we filtered each subnetwork according to the differential interactome of the respective disorder-manifesting tissue, which was calculated using the *diff_i* method. We included only nodes adjacent to differential interactions scoring at the top 1% within the respective tissue interactome (Fig. 3A, right panel). The top 1% filtered subnetworks were much smaller than the unfiltered subnetworks, with a median of 34 versus 3,000 genes, respectively. Next, we tested whether filtered subnetworks were more enriched for additional causal genes relative to the unfiltered subnetworks (see Methods). We collated causal genes according to tissues. For the 10 tissues with more than 30 genes, we compared between the percentage of causal genes in the filtered and unfiltered subnetworks using paired Wilcoxon test. In 5/10 tissues, filtered subnetworks were significantly more enriched (p≤0.05) (Fig. 3B). Next, we tested for enrichment in each individual case. In 283 of the 1,527 cases (18.6%), the filtered subnetwork was enriched significantly for additional causal genes (p≤0.05, Fisher exact test, adjusted via Benjamini-Hochberg procedure). This success rate was not uniform across tissues, with some tissues showing high rates (e.g., heart 53%) and other tissues showing lower rates (e.g., testis 9.5%; Table S5).To find if this success rate was expected by chance, we permuted the disease-tissue associations and repeated this analysis 1,000 times (see Methods), showing that this success rate was highly statistically significant (p≤0.001, Fig. 3C). We further tested how the enrichment of the disorder-manifesting tissue interactome compared to the enrichments in other tissue interactomes. For that, we compared the significance of the enrichment in the ‘accurate’ tissue to the median significance of the enrichments in other tissues, using Mann-Whitney-U test between the p-value in the disease tissue and the median p-value across all tissues. This confirmed that the p-value in the disease tissue is significantly lower then the p-values in other tissues (p = 6.19e-68).

**Fig. 3.**
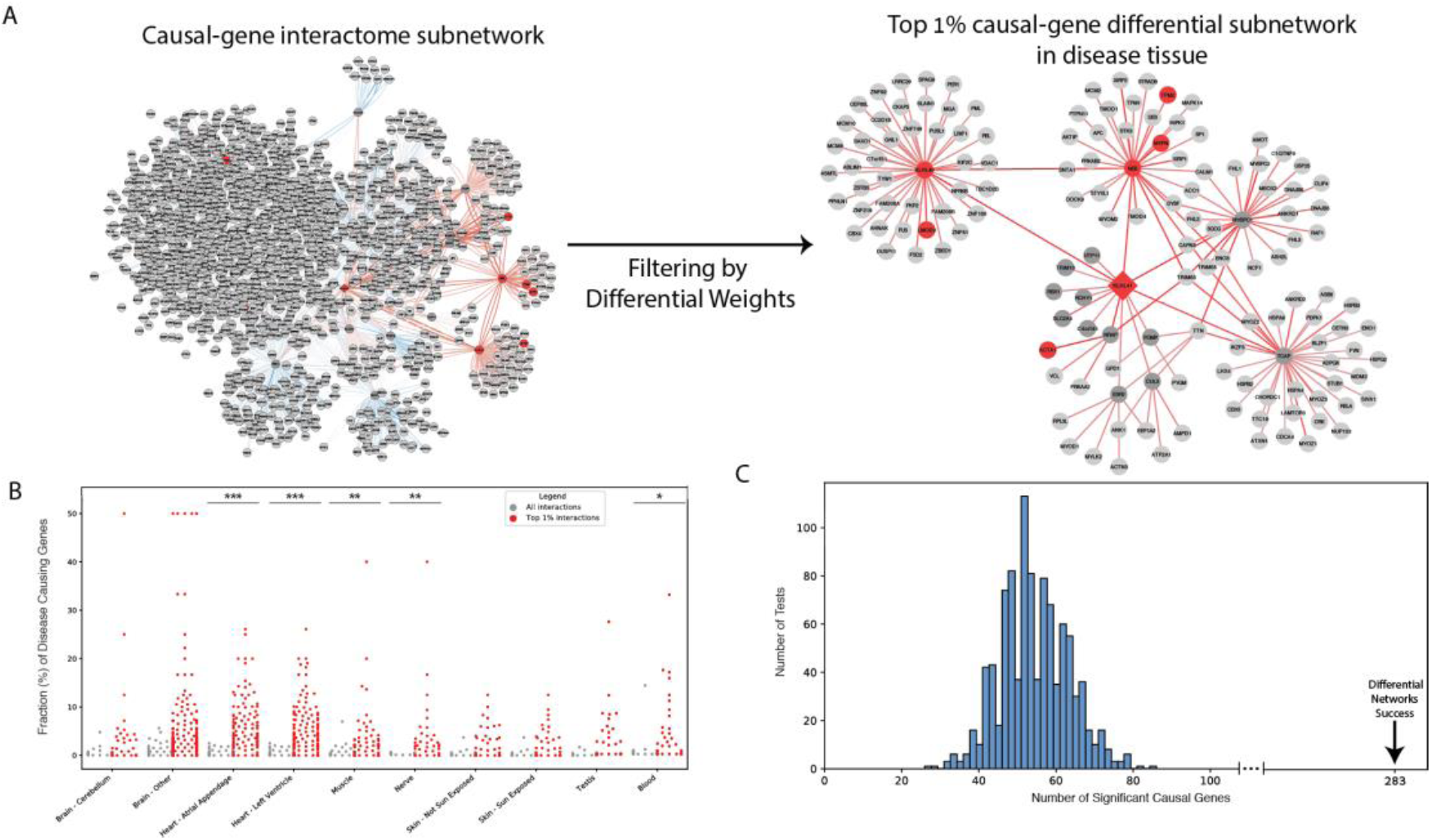
Assessing the ability of differential tissue interactomes to uncover disorder-related genes. A. Left: The interactome subnetwork surrounding the query gene KLH41 that is causal for spinocerebellar ataxia included its direct and secondary interactions. The subnetwork amounted to 1,745 genes, including 8 other genes causal for nemaline myopathy. Right: The top 1% filtered subnetwork of KLH41 in skeletal muscle, where nemaline myopathy is manifested. The subnetwork contained 144 genes, including 6/8 of the other genes causal for nemaline myopathy (p=6.97E-06). Genes causal for nemaline myopathy appear in red; KLH41 is shaped as a diamond. B. Visualization of the tissue-based enrichment analysis. Each dot represents a query gene in the unfiltered subnetwork (grey) and the top 1% filtered subnetwork of its disease-manifesting tissue (red). The value of each dot corresponds to the percentage of genes causal for the same disease as the query gene, out of the total number of nodes in that subnetwork. Data are shown for the 10 tissues with at least 30 different causal genes. In 5/10 tissues the top 1% filtered subnetworks were significantly enriched (paired Wilcoxon test; * p≤ 0.05, ** p≤ 5E-3, *** p≤ 1E-17). C. The distribution of the numbers of successes in the 1,000 randomization runs, showing a median of 54 successes and maximal value of 86. In contrast, the tested *diff_i* method had 283 successes (p≤0.001) which places it far beyond any of the randomized test.

The power of this analysis is demonstrated by the case of KLH41 (Fig. 3A). This protein is involved in skeletal muscle development and maintenance processes. Mutations in this gene cause nemaline myopathy (NM), a rare hereditary muscle disorder. The interactome subnetwork of KLH41 included 1,745 nodes, among which were eight other genes causal for NM. Upon filtering this subnetwork to include only nodes with differential interactions scoring at the top 1%, we obtained a much smaller subnetwork containing only 144 genes, yet, including six of the eight additional causal genes (Fisher exact test p=6.97E-06, Fig. 3A). Notably, in the case of KLH41, some of the causal genes were only indirectly connected to each other, showing the ability of differential interactomes to pull out meaningful secondary relationships. Additional examples appear in Fig. 4. Thus, differential tissue interactomes appear to be effective in illuminating disorder mechanisms.

**Fig. 4.**
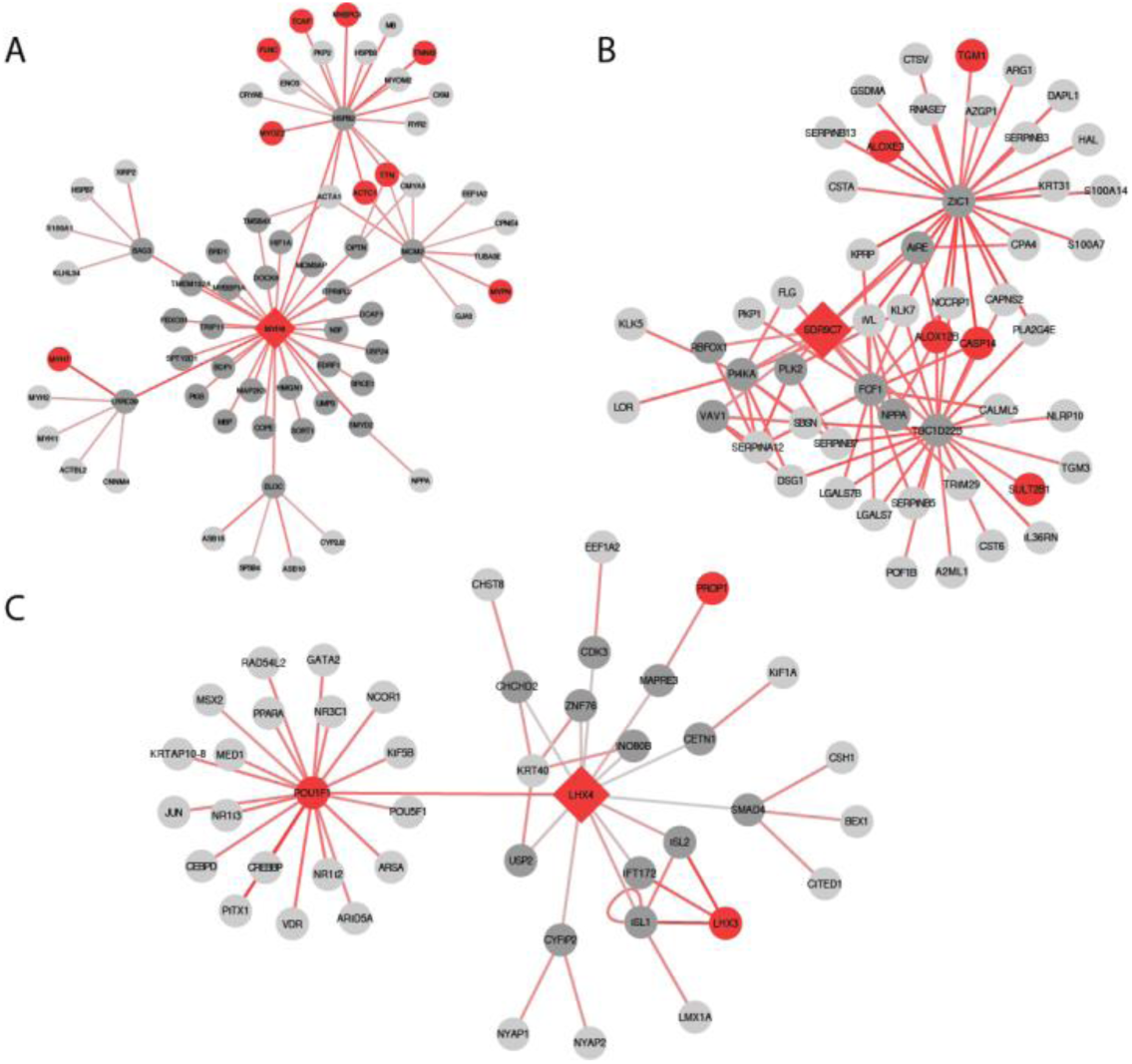
The top 1% differential subnetwork that surrounds a causal gene (red diamond) in its disorder-tissue interactomes is enriched for additional causal genes. A. MYH6 is casual for familial hypertrophic cardiomyopathy. Its filtered subnetwork in heart contained 64 genes (compared to 2,277 in the unfiltered subnetwork), including all 9 genes casual for hypertrophic cardiomyopathy in the unfiltered subnetwork (p=6.17E-15). B. LHX4 is casual for pituitary hormone deficiency. Its filtered subnetwork in the pituitary contained 44 genes (compared to 1,800 genes in the unfiltered subnetwork), including all 3 genes causal for this disorder in the unfiltered subnetwork (p= 1.31E-05). C. SDR9C7 is casual for autosomal recessive congenital Ichthyosis that manifests in skin. Its filtered subnetwork in skin contained 40 genes (compared to 652 genes), including 4 of the 5 genes causal for this disorder in the unfiltered subnetwork (p=5.85E-05).

## DISCUSSION

Differential network analysis is a powerful paradigm in network biology (Ideker and Krogan, 2012), but has not been widely applied to tissue interactomes composed of PPIs. This is partly because there were no large-scale quantitative data allowing for interactome weighting in different contexts, and because benchmarks allowing for evaluation of different schemes have been lacking. Here we show rigorously and at large-scale the value of differential network analysis in highlighting tissue-specific processes and disease mechanisms. To allow for large-scale assessments, we created two manually-curated datasets. The first dataset includes 7,718 associates between 6,499 GO biological process terms and 48 tissues. The second dataset includes 1,527 associations between 1,185 hereditary disorders and 26 clinically-manifesting tissues (Fig. 1). While these resources remain incomplete and are not distributed uniformly across tissues, they compose a uniquely extensive resource that may serve many other future studies and applications focusing on the physiology and pathophysiology of human tissues.

We implemented and assessed five different interactome weighting methods, including four DN methods. We first analyzed the ability of each method to highlight tissue-specific processes. The non-differential method of ranking genes by their expression within a tissue (*expr_n*), was outperformed by DN methods, demonstrating the value of differential weighting schemes in highlighting tissue-specific features (Fig. 2A). Both node-based and interaction-based DN methods performed well in highlighting tissue-specific processes (Fig. 2C). The value of interaction-based methods was also assessed using the gene-neighbors test (Fig. 2E). Given that interactomes contain more interactions than nodes, scoring by interactions might be more informative in some settings.

After identifying the power of the DN methods and establishing that tissue-specific processes are well captured by focusing on differential interactions, we turned to analyze hereditary disorders. Hereditary disorders tend to manifest clinically in few tissues, while the aberrant gene is present and often expressed across the body (Fig. 1D) (Lage, 2014; Barshir *et al.*, 2014, 2018). However, the mechanisms underlying the tissue-selective manifestation of hereditary disorders is well understood only in few cases; for most disorders it remains enigmatic (Barshir *et al.*, 2014). Previous efforts to unravel the molecular basis of this phenomenon showed that in a considerable fraction of hereditary disorders, clinically affected tissues are associated with elevated expression of the causal gene (Barshir *et al.*, 2014; Lage, 2014), tissue-specific interactions (Barshir *et al.*, 2014), and down-regulation of paralogs of the casual genes correct this ref (Barbeira *et al.*, 2018). These properties easily translate into differential nodes and interactions within tissue interactomes, suggesting that differential subnetworks surrounding causal genes could be informative of additional genes causal for the same disorder.

To answer this challenge, we assessed whether differential network analysis can effectively identify disorder-related genes. For this, we focused only on the top 1% differential interactions surrounding causal genes. This stringent cutoff reduced subnetworks sizes from ~3,000 genes to ~30 genes (Fig. 3A,D). In half of the tissues tested, the differential subnetworks were enriched significantly for disorder-related genes, relative to unfiltered subnetworks. In 18.6% of the individual cases, enrichment was highly significant, and the fraction of successful cases was much higher than expected by chance (p≤0.001). With the rapid accumulation of molecular interactions data and the move towards precision medicine, differential and other context-sensitive filtering approaches are becoming essential for meaningful interactome analyses and interpretation (Gligorijević *et al.*, 2016). The methods we described, the large-scale resources, and the rigorous assessment tests that we put presented, may thus serve many other future applications.

## METHODS

### Tissue expression data

RNA sequencing profiles were obtained from the GTEx portal (version 7) (GTEx Consortium, 2017), resulting in 11,216 samples from 51 tissues. Only genes with more than 5 read counts in at least 10 samples were included in the analysis. Raw read counts were normalized for sample library size via the TMM method by edgeR (Robinson *et al.*, 2010) to produce counts per million (cpm). Only genes with cpm values ≥8 in at least 10 samples were considered henceforth, and their cpm values were log2 transformed to obtain normal distributions. Samples per tissue were merged such that the expression of each gene was set to its median expression value across samples. Similarly to (Paulson *et al.*, 2017), brain sub-regions were further merged into three regions, named basal ganglia, cerebellum, and ‘other’.

### PPI data

Human PPIs were gathered from BioGrid (Chatr-Aryamontri *et al.*, 2017), DIP (Salwinski *et al.*, 2004), MINT (Ceol *et al.*, 2010) and IntAct (Aranda *et al.*, 2010) by using the MyProteinNet web-server (Basha *et al.*, 2015). The MyProteinNet web-server ensures that only PPIs detected by well-established methods for physical interactions detection were considered.

### Construction of tissue interactomes

All tissue interactomes contained the same number of nodes and interactions and differed only in the weights that they were associated with. We used five different weighting methods, as described below. Let 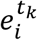 denote the normalized read counts of gene *i* in tissue *t*_*k*_.

1. The non-differential method, *expr_n:* In the interactome of tissue *t*_*k*_, each node was assigned with a weight that reflects its expression level in that tissue, 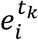.
2. The differential node method, *diff_n*: In the interactome of tissue *t*_*k*_, each node was assigned with a weight, denoted 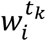, that reflects its differential expression level in that tissue relative to its median expression level across all tissues (equation 1):

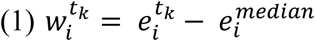
3. The differential node method, *pref_n*: In the interactome of tissue *t*_*k*_, each node was assigned with a weight, denoted 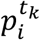, that reflects its preferential expression in that tissue as computed by (Sonawane *et al.*, 2017).
4. The differential interaction method, *pref_i*: In the interactome of tissue *t*_*k*_, an interaction between genes *i* and *j* was assigned a weight, denoted 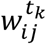, that was set tothe sum of the preferential weight of the two nodes (equation 2):

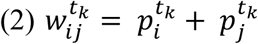
5. The differential interaction method, *diff_i*: In the interactome of tissue *t*_*k*_, an interaction between genes *i* and *j* was assigned a weight, denoted 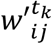, that was (Basha, Shpringer, *et al.*, 2017). This was approximated by 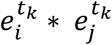, namely the sum of their log2 normalized counts values (equation 3). We further normalized this weight relative to the maximal interaction weight in that tissue, to fit the range of [0,1] (equation 4). The differential weight of that interaction, denoted 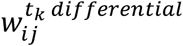, was computed by subtracting the median weight for that interaction across all tissues (equation 5).

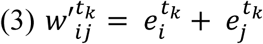

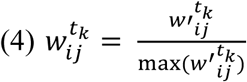

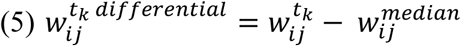

### Associating GO terms with tissues

We created a list of tissue-related keywords; for example, heart-related keywords included ‘myo’ and ‘cardiac’ (full list is in Table S4). We associated GO biological process terms to tissues by searching the GO term names for tissue-related keywords. We included associations of GO term to tissues created by (Greene *et al.*, 2015), after mapping BRENDA tissues to GTEx tissues. Lastly, we manual checked each GO term – tissue association (Fig. 1A,B, Table S1).

### Interactomes GO enrichment tests

All GO enrichments were conducted in GOrilla (Eden *et al.*, 2009) using the ranked list option. The GOrilla web-server calculates GO enrichment for a ranked list by taking the optimal hypergeometric tail probability that is found over all possible partitions induced by the gene ranking and corrected for multiple hypothesis testing (Eden *et al.*, 2007). Only processes with p-values lower than 10^-3^ were considered enriched. To test the GO enrichment of a tissue interactome, we converted each interactome into a ranked list of genes. For node-based methods, the list was ordered in descending order according to node weights. For interaction-based methods, we assigned each node its median weight across all of its interactions in the interactome, and the list was ordered in descending order according to these weights. We repeated this analysis upon assigning each node with its maximal interaction weight, but results were inferior compared to the median (data not shown).

### Enrichment specificity tests

To determine if a ranked list was enriched specifically for tissue-specific GO biological process terms we used Fisher’s exact test. For this, we assessed the overlap between the GO terms associated with the tissue, and the GO terms associated with any tissue that were enriched in the ranked list (Fig. 2B). Tests with a p-value ≤ 0.05 were considered successful.

### Differential interactions tests

GO terms and associations were gathered from the MyGene.info web-service (Xin *et al.*, 2015) (Sep 2018). We considered only genes with (i) manually curated tissue-associated GO biological process terms, (ii) ≥ 5 interactors related to any of these tissue-associated GO terms, and (iii) ≥ 5 interactors that were not related to any of these tissue-associated GO terms. For each such query gene, we divided its interactions into two groups: Group A contained interactions with interactors that were annotated to a tissue-associated GO term as the query gene itself, and group B contained all its other interactions. Interactions weights were calculated using the *diff_i* method. To test the null hypothesis that the weights of interactions in group A were not significantly higher than the weights of interactions in group B, we performed the Mann-Whitney-U test. To check for a general trend per tissue, we collapsed together all the interactions for the same query gene, by computing the median of each A and B group. We applied this procedure to query genes belonging to tissues with ≥ 20 query genes. To test for a statistically significant difference between the groups per tissue, we applied paired Wilcoxon test to the two lists of medians per tissue (Fig. 2E). All p-values were adjusted for multiple hypothesis testing by using Benjamini-Hochberg procedure.

### Genetic disorders and causal genes datasets

Data of genetic disorders and causal genes with known molecular basis were downloaded from OMIM on January 2018 (Amberger *et al.*, 2015). To associate genetic disorders with tissues we used the phenotypic series data from OMIM. A phenotypic series is an aggregation of genetic disorders with a common phenotype. We associated each phenotypic series with its clinically manifesting tissues (Fig. 1A,C) by manually reviewing the information in OMIM (disorder description and clinical phenotypes) and other sources, such as NIH Genetic Home Reference (National Library of Medicine, 2013) and the Genetic and Rare Disease Information Center (GARD) (Lewis *et al.*, 2017). A phenotypic series could be classified to more than one tissue. Causal genes for similar disorders were considered related (Table S3). Disorder-associated tissues were manually matched with relevant GTEx tissues.

### Differential interactome enrichment for disorder-related genes

We define a disorder case as a tuple (disorder; causal gene; manifesting tissue). For each case, we considered an interactome subnetwork composing the direct and secondary interactions of the casual gene (denoted query gene). We limited our analyses to causal genes whose interactome subnetwork contained other genes causal for a similar disorder. We filtered each subnetwork, such that it contained only nodes with differential interactions that weighted at the top 1% according to the *diff_i* method. We calculated the significance of the enrichment of the filtered subnetwork for genes causal for a similar disorder as the query gene, by using Fisher exact test with p-values adjusted for multiple hypothesis testing by Benjamini-Hochberg procedure. We then assessed the overlap of disorder causal genes between the nodes connected by interactions in top 1% weights and nodes connected by less weighted interactions. Cases with adjusted p-value ≤ 0.05 were considered significantly enriched.

To test the null hypothesis that enrichments in the disorder-manifesting tissue was not more significant than enrichments in other tissues, we compared between the enrichment p-value obtained in the disorder-manifesting tissue and the median enrichment p-value across all tissues, using the Mann-Whitney-U test.

To check the trend per tissue, we analyzed tissues with over 30 cases. For each tissue we two groups: (A) the percentage of disease genes in the unfiltered subnetwork and (B) the percentage of disease genes in the filtered subnetwork. We performed a Paired-Wilcoxon test between groups A and B, to test the null hypothesis that the fraction of the causal genes in the unfiltered subnetwork (group A) does not differ from the fraction of causal genes in the filtered subnetwork (group B).

To test the null hypothesis that the number of filtered subnetworks that were enriched significantly (p ≤ 0.05), denoted *num_s*, was not higher than expected by chance, we carried randomization tests. In each test, for each causal gene we selected a tissue at random and tested whether the filtered subnetwork of the randomly selected tissue interactome was enriched significantly for additional causal genes. We repeated this for each causal gene and recorded the number of significantly enriched random subnetworks, *num_r*. We repeated this procedure 1,000 times. The significance of *num_s* was calculated as the fraction of randomized runs with *num_r* ≥ *num_s*.

## AWKNOLEDGEMENTS

We thank the members of the Yeger-Lotem lab for their support in this research.

## REFERENCES

Amberger, J.S. et al. (2015) OMIM.org: Online Mendelian Inheritance in Man (OMIM??), an Online catalog of human genes and genetic disorders. Nucleic Acids Res., 43, D789–D798.

Aranda, B. et al. (2010) The IntAct molecular interaction database in 2010. Nucleic Acids Res, 38, D525–31.

Bandyopadhyay, S. et al. (2010) Rewiring of genetic networks in response to DNA damage. Science (80-.)., 330, 1385–1389.

Barbeira, A.N. et al. (2018) Exploring the phenotypic consequences of tissue specific gene expression variation inferred from GWAS summary statistics. Nat. Commun., 9.

Barshir, R. et al. (2014) Comparative analysis of human tissue interactomes reveals factors leading to tissue-specific manifestation of hereditary diseases. PLoS Comput Biol, 10, e1003632.

Barshir, R. et al. (2018) Role of duplicate genes in determining the tissue-selectivity of hereditary diseases. PLoS Genet.

Basha, O. et al. (2015) MyProteinNet: Build up-to-date protein interaction networks for organisms, tissues and user-defined contexts. Nucleic Acids Res., 43.

Basha, O., Shpringer, R., et al. (2017) The DifferentialNet database of differential protein– protein interactions in human tissues. Nucleic Acids Res.

Basha, O., Barshir, R., et al. (2017) The TissueNet v.2 database: A quantitative view of protein-protein interactions across human tissues. Nucleic Acids Res., 45, D427–D431.

Ceol, A. et al. (2010) MINT, the molecular interaction database: 2009 update. Nucleic Acids Res, 38, D532–9.

Chatr-Aryamontri, A. et al. (2017) The BioGRID interaction database: 2017 update. Nucleic Acids Res., 45, D369–D379.

Eden, E. et al. (2007) Discovering motifs in ranked lists of DNA sequences. PLoS Comput. Biol.

Eden, E. et al. (2009) GOrilla: a tool for discovery and visualization of enriched GO terms in ranked gene lists. BMC Bioinformatics, 10, 48.

Gambardella, G. et al. (2013) Differential network analysis for the identification of condition-specific pathway activity and regulation. Bioinformatics, 29, 1776–1785.

Gill, R. et al. (2010) A statistical framework for differential network analysis from microarray data. BMC Bioinformatics, 11, 95.

Gligorijević, V. et al. (2016) Integrative methods for analyzing big data in precision medicine. Proteomics.

Goenawan, I.H. et al. (2016) DyNet: Visualization and analysis of dynamic molecular interaction networks. In, Bioinformatics., pp. 2713–2715.

Greene, C.S. et al. (2015) Understanding multicellular function and disease with human tissue-specific networks. Nat Genet, 47, 569–576.

GTEx Consortium (2017) Genetic effects on gene expression across human tissues. Nature, 550, 204–213.

Ha, M.J. et al. (2014) DINGO: Differential network analysis in genomics. Bioinformatics, 31, 3413–3420.

Ideker, T. and Krogan, N.J. (2012) Differential network biology. Mol. Syst. Biol., 8.

Islam, M.F. et al. (2013) Comparative analysis of differential network modularity in tissue specific normal and cancer protein interaction networks. J. Clin. Bioinforma., 3, 19.

Kitsak, M. et al. (2016) Tissue Specificity of Human Disease Module. Sci. Rep.

Lage, K. et al. (2008) A large-scale analysis of tissue-specific pathology and gene expression of human disease genes and complexes. Proc. Natl. Acad. Sci. U. S. A., 105, 20870–5.

Lage, K. (2014) Protein-protein interactions and genetic diseases: The interactome. Biochim. Biophys. Acta - Mol. Basis Dis., 1842, 1971–1980.

Landeghem, S. Van et al. (2016) Diffany: an ontology-driven framework to infer, visualise and analyse differential molecular networks. BMC Bioinformatics, 17, 18.

Lewis, J. et al. (2017) Marking 15 years of the Genetic and Rare Diseases Information Center. Transl. Sci. Rare Dis.

National Library of Medicine. (US). G.H.R. [Internet]. B. (MD): T. (2013) Genetic Home Reference.

Luck, K. et al. (2017) Proteome-Scale Human Interactomics. Trends Biochem. Sci., 42, 342–354.

Ma, C. et al. (2014) Machine Learning-Based Differential Network Analysis: A Study of Stress-Responsive Transcriptomes in Arabidopsis. Plant Cell, 26, 520–537.

Magger, O. et al. (2012) Enhancing the prioritization of disease-causing genes through tissue specific protein interaction networks. PLoS Comput Biol, 8, e1002690.

Marbach, D. et al. (2016) Tissue-specific regulatory circuits reveal variable modular perturbations across complex diseases. Nat. Methods, 13, 366–370.

Paulson, J.N. et al. (2017) Tissue-aware RNA-Seq processing and normalization for heterogeneous and sparse data. BMC Bioinformatics, 18.

Pierson, E. et al. (2015) Sharing and Specificity of Co-expression Networks across 35 Human Tissues. PLoS Comput. Biol.

Regev, A. et al. (2017) Science Forum: The Human Cell Atlas. Elife.

Robinson, M.D. et al. (2010) edgeR: a Bioconductor package for differential expression analysis of digital gene expression data. Bioinformatics, 26, 139–140.

Salwinski, L. et al. (2004) The Database of Interacting Proteins: 2004 update. Nucleic Acids Res, 32, D449–51.

Sonawane, A.R. et al. (2017) Understanding Tissue-Specific Gene Regulation. Cell Rep., 21, 1077–1088.

Uhlen, M. et al. (2015) Tissue-based map of the human proteome. Science (80-.)., 347, 1260419–1260419.

Warsow, G. et al. (2013) Differential network analysis applied to preoperative breast cancer chemotherapy response. PLoS One, 8.

Xin, J. et al. (2015) MyGene.info and MyVariant.info: Gene and Variant Annotation Query Services.

Yao, V. et al. (2018) Enabling Precision Medicine through Integrative Network Models. J. Mol. Biol.

Yeger-Lotem, E. and Sharan, R. (2015) Human protein interaction networks across tissues and diseases. Front Genet, 6, 257.

Zickenrott, S. et al. (2016) Prediction of disease-gene-drug relationships following a differential network analysis. Cell Death Dis., 7, e2040.

