## Supplementary material for "Differential network analysis of human tissue interactomes highlights tissue-selective processes and genetic disorder genes": Fig S1; Table S1; Table S2; Table S3; Table S4; Table S5

### Supporting information

**Figure S1. Weight distribution histograms for the different weighting schemes.**

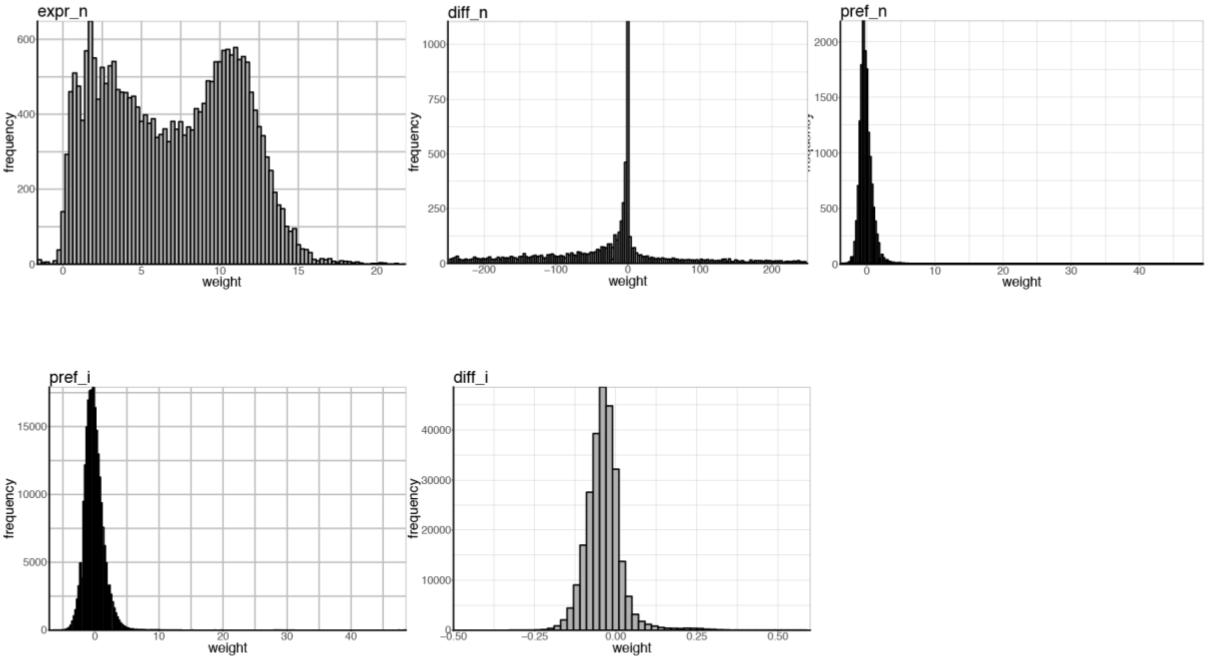


**Table S1. Manually curated tissue-associated GO terms.** Due to its large size, the table is accessible in tsv format via this link: http://netbio.bgu.ac.il/diffnet_files/TableS1.csv

**Table S2. The enrichment specificity test for the different weighting approaches.** The table shows the p-values obtained via Fisher's exact test per tissue. Significant p-values ($p \leq 0.05$) appear in red.

| **Tissue** | ***expr_n*** | ***diff_n*** | ***pref_n*** | ***pref_i*** | ***diff_i*** |
|---|---|---|---|---|---|
| Adipose - Subcutaneous | 1 | 2.61E-03 | 1.28E-02 | 1.59E-01 | 9.73E-02 |
| Adipose - Visceral (Omentum) | 1 | 1.09E-02 | 3.25E-03 | 3.43E-01 | 1.92E-01 |
| Adrenal Gland | 1 | 1 | 1 | 1 | 1.93E-02 |
| Artery – Aorta | 1 | 6.41E-01 | 2.76E-01 | 5.57E-01 | 6.76E-01 |
| Artery – Coronary | 7.72E-01 | 7.10E-01 | 5.55E-02 | 3.03E-02 | 6.28E-01 |
| Artery – Tibial | 8.00E-01 | 6.56E-01 | 2.76E-01 | 2.91E-01 | 6.52E-01 |
| Brain - Basal Ganglia | 1.06E-19 | 1.06E-45 | 1.77E-37 | 2.74E-36 | 1.09E-48 |
| Brain – Cerebellum | 1.15E-31 | 6.08E-42 | 2.76E-32 | 1.07E-33 | 6.56E-44 |
| Brain – Other | 1.86E-23 | 1.67E-48 | 1.32E-35 | 2.55E-33 | 6.84E-40 |
| Breast - Mammary Tissue | 7.41E-01 | 8.74E-01 | 1 | 1 | 7.56E-01 |
| Colon – Sigmoid | 1 | 1 | 1 | 1 | 5.37E-01 |
| Colon – Transverse | 1 | 1 | 1 | 1 | 1 |
| Esophagus - Gastroesophageal Junction | 1 | 1 | 1 | 1 | 1 |
| Esophagus – Mucosa | 1 | 1 | 1 | 1 | 1 |
| Esophagus - Muscularis | 1 | 1 | 1 | 1 | 9.19E-02 |
| Heart - Atrial Appendage | 3.37E-10 | 8.10E-29 | 9.40E-40 | 1.13E-25 | 1.66E-31 |
| Heart - Left Ventricle | 1.56E-12 | 2.01E-24 | 3.50E-39 | 4.07E-26 | 5.02E-38 |
| Kidney – Cortex | 6.40E-02 | 1.38E-29 | 3.57E-38 | 1.40E-29 | 3.83E-39 |
| Liver | 5.39E-01 | 1.97E-01 | 3.96E-01 | 3.90E-01 | 1.70E-01 |
| Lung | 8.34E-01 | 9.66E-01 | 1 | 1 | 1.69E-01 |
| Minor Salivary Gland | 1 | 3.62E-01 | 1.43E-01 | 1.37E-01 | 3.33E-01 |
| Muscle – Skeletal | 4.42E-40 | 1.33E-50 | 1.86E-64 | 3.78E-62 | 1.43E-69 |
| Nerve – Tibial | 6.01E-02 | 3.99E-04 | 1.19E-05 | 5.42E-03 | 2.56E-14 |
| Ovary | 1 | 5.21E-01 | 4.94E-04 | 1.91E-04 | 2.16E-06 |
| Pancreas | 1 | 1.43E-01 | 2.50E-16 | 2.50E-16 | 1.07E-10 |
| Pituitary | 1 | 1 | 1 | 1 | 2.66E-01 |
| Prostate | 1 | 4.47E-02 | 8.30E-06 | 6.67E-06 | 7.60E-04 |
| Skin - Sun Exposed (Lower leg) | 2.13E-06 | 8.67E-16 | 3.20E-25 | 1.95E-22 | 3.60E-23 |
| Small Intestine - Terminal Ileum | 2.19E-01 | 3.12E-01 | 1.42E-05 | 2.81E-05 | 2.35E-04 |
| Stomach | 1 | 2.25E-01 | 1.55E-02 | 7.33E-03 | 3.64E-03 |
| Testis | 7.23E-10 | 4.17E-28 | 5.72E-15 | 1.34E-13 | 3.79E-18 |
| Thyroid | 7.12E-06 | 3.50E-05 | 7.91E-15 | 9.98E-11 | 4.70E-09 |
| Uterus | 5.85E-01 | 7.81E-01 | 1.91E-01 | 1.04E-01 | 3.47E-01 |
| Whole Blood | 3.09E-67 | 5.37E-81 | 4.14E-85 | 2.37E-93 | 3.48E-114 |
| # of significant tissues | 10 | 15 | 19 | 18 | 18 |

**Table S3. Manually curated tissue-associated hereditary diseases and causal genes.** Due to its large size, the table is accessible in tsv format via this link: http://netbio.bgu.ac.il/diffnet_files/TableS3.tsv

**Table S4. Tissue-related keywords**.

| Searched item | GTEx tissue |
|---|---|
| adipo | Adipose |
| fat | Adipose |
| adrenal | Adrenal Gland |
| adreno | Adrenal Gland |
| pheochromocytoma | Adrenal Gland |
| arter | Artery |
| accumbens | Brain |
| amygdala | Brain |
| basal ganglia | Brain |
| brain | Brain |
| caudate | Brain |
| cerebel | Brain |
| cerebr | Brain |
| glia | Brain |
| hippocamp | Brain |
| putamen | Brain |
| substantia nigra | Brain |
| thalam | Brain |
| breast | Breast |
| mammar | Breast |
| appendix | Colon |
| colon | Colon |
| gastr | Colon |
| recta | Colon |
| rectum | Colon |
| sigmoid | Colon |
| transvers | Colon |
| esophag | Esophagus |
| card | Heart |
| cardiac muscle | Heart |
| heart | Heart |
| kidney | Kidney |
| renal | Kidney |
| reno | Kidney |
| biliary vesicle | Liver |
| hepat | Liver |

| | |
|---|---|
| liver | Liver |
| lung | Lung |
| pulmo | Lung |
| salivary gland | Minor Salivary Gland |
| salivarygland | Minor Salivary Gland |
| muscl | Muscle |
| muscle | Muscle |
| muscul | Muscle |
| myo | Muscle |
| skeletal | Muscle |
| skeletal muscle | Muscle |
| skeletalmuscle | Muscle |
| smooth | Muscle |
| smooth muscle | Muscle |
| smooth muscle | Muscle |
| striated | Muscle |
| gonad | Ovary |
| ovar | Ovary |
| ovarian tube | Ovary |
| pancrea | Pancreas |
| pituitar | Pituitary |
| prostat | Prostate |
| skin | Skin |
| duoden | Small Intestine |
| gastr | Small Intestine |
| ileo | Small Intestine |
| ileum | Small Intestine |
| intestin | Small Intestine |
| jejun | Small Intestine |
| smallintestine | Small Intestine |
| entero | Stomach |
| gastr | Stomach |
| stomach | Stomach |
| gonad | Testis |
| testes | Testis |
| testis | Testis |
| thyroid | Thyroid |
| placent | Uterus |
| uterin | Uterus |
| uterine tube | Uterus |

**Table S5**. **Success of predicting disease genes per tissue.**

| Tissue | Number of cases | Successes (Fisher exact test p ≤ 0.05 after BH procedure) | Percentile of success |
|---|---|---|---|
| Heart - Atrial Appendage | 142 | 76 | 53.52 |
| Heart - Left Ventricle | 160 | 84 | 52.5 |
| Muscle - Skeletal | 89 | 25 | 28.08 |
| Nerve - Tibial | 48 | 8 | 16.66 |
| Artery - Aorta | 19 | 3 | 15.78 |
| Lung | 28 | 4 | 14.28 |
| Esophagus - Mucosa | 8 | 1 | 12.5 |
| Whole Blood | 52 | 6 | 11.53 |
| Skin - Not Sun Exposed (Suprapubic) | 95 | 10 | 10.52 |
| Skin - Sun Exposed (Lower leg) | 95 | 9 | 9.47 |
| Testis | 74 | 7 | 9.45 |
| Brain - Other | 511 | 47 | 9.19 |
| Pituitary | 20 | 1 | 5 |
| Brain - Cerebellum | 64 | 2 | 3.12 |
| Ovary | 23 | 0 | 0 |
| Liver | 18 | 0 | 0 |
| Artery - Coronary | 12 | 0 | 0 |
| Artery - Tibial | 12 | 0 | 0 |
| Colon - Sigmoid | 12 | 0 | 0 |
| Thyroid | 10 | 0 | 0 |
| Brain – Basal Ganglia | 9 | 0 | 0 |
| Adipose - Subcutaneous | 6 | 0 | 0 |
| Adipose - Visceral (Omentum) | 6 | 0 | 0 |
| Breast | 5 | 0 | 0 |
| Uterus | 5 | 0 | 0 |
| Esophagus - Gastroesophageal Junction | 4 | 0 | 0 |
| Total | 1527 | 283 | 18.53 |
